# Large-Scale In Vivo Electrophysiology and Dimensionality-Reduction Analysis Reveal Opposing Thalamic and Hippocampal Excitability After Third-Trimester–Equivalent Alcohol Exposure in Mice

**DOI:** 10.1101/2025.06.02.657410

**Authors:** M.D. Morningstar, G. Acosta, S.A. McKenzie, C.F. Valenzuela, B.J. Clark, D. N. Linsenbardt

## Abstract

Acute, binge-like alcohol exposure during the third-trimester–equivalent period (TTAE) produces apoptotic neurodegeneration in brain regions critical for spatial learning and memory, including the anterior thalamic nuclei (ATN), which are necessary for contextual learning and memory. To investigate the neural mechanisms underlying spatial cognition deficits following developmental alcohol exposure, we performed high-density electrophysiological recordings in and around the ATN of behaving mice, sampling 7,490 neurons across multiple, predominantly thalamic depths. To model acute binge-like TTAE, C57BL/6J mice received two injections of 2.5 g/kg ethanol (or saline) on postnatal day (PND) 7. In young adulthood (>PND 60), mice were implanted with silicon or multi-wire electrode arrays and allowed to explore a circular arena (40 cm diameter) under dim red light with two wall-mounted LED cues that rotated pseudo-randomly. Spike timing and waveform features were extracted from each putative single unit. Across the recorded population, firing rates were generally reduced following TTAE, although responses were heterogeneous. To characterize this variability, neuronal features were embedded using uniform manifold approximation and projection (UMAP). Logistic regression classified saline- and TTAE-exposed neurons based on electrophysiological properties and UMAP coordinates, achieving 80% accuracy. Cluster-based comparisons showed weak correspondence between physiological similarity and treatment response, suggesting TTAE effects are driven primarily by anatomical location. This study presents the first large-scale *in vivo* electrophysiological recordings in a freely behaving mouse model of fetal alcohol spectrum disorders, revealing divergent circuit adaptations marked by hippocampal hyperexcitability and thalamic dysfunction.

## Introduction

Fetal Alcohol Spectrum Disorders (FASDs) encompass a range of neurodevelopmental abnormalities resulting from in utero alcohol exposure. These conditions are associated with lasting cognitive, behavioral, and neurological impairments, including deficits in spatial learning and memory. One particularly vulnerable developmental window is the third trimester of human pregnancy, which in rodents corresponds to the early postnatal period ^1^. In some regions of the United States, third-trimester alcohol exposure (TTAE) accounts for up to 8% of FASD cases ^2^.

Emerging evidence indicates that the hippocampal-diencephalic-cingulate (HDC) network is a key target of TTAE. First described by Papez and later refined through cross-species research, this circuit includes the hippocampal formation (HPF), mammillary bodies, retrosplenial cortex (RSC), and anterior thalamic nuclei (ATN) ^3–5^. The HDC network enables allocentric navigation, contextual learning, and the integration of spatial cues.

Within this network, neurons in the ATN serve key roles in contextual memory and learning. These neurons encode directional information using both internal cues (e.g., vestibular, proprioceptive, and motor signals) and external landmarks ^6,7^. Their activity is shaped by inhibitory input from the thalamic reticular nucleus (TRN) and modulated by reciprocal connections with the RSC and HPF ^8–10^. Disruption of this circuitry through lesions to the ATN or RSC leads to impaired navigation, underscoring the importance of bidirectional thalamocortical communication in spatial cognition.

Acute, binge-like alcohol exposure on postnatal day (PND)7 in rodent models induces widespread neuronal apoptosis in components of the HDC network, including the subiculum, RSC, and ATN ^11,12^. In humans, prenatal alcohol exposure, likely involving the third trimester, has been linked to thalamic volume loss and reduced activation during spatial memory tasks ^13,14^. Additional rodent studies show long-term morphological changes in thalamocortical pathways and sex-dependent alterations in functional connectivity ^15,16^. The anterodorsal nucleus, in particular, shows functional impairments following TTAE, including decreased intrinsic excitability, diminished excitatory postsynaptic currents in RSC-projecting AD neurons ^17^ and reduced Fos expression following contextual learning ^18^, suggesting broad circuit-level dysfunction.

Despite well-documented anatomical and molecular disruptions following acute binge-like TTAE, its functional impact on ATN activity during behavior remains poorly understood, particularly in awake, freely moving animals. We hypothesized that acute TTAE produces long-lasting alterations in the functional properties of ATN neurons that are detectable *in vivo* during naturalistic behavior. To test this, we used high-density electrophysiological recordings to examine neural activity in and around the ATN of behaving mice, compiling a total of 7490 neurons across multiple recording days and depths.

We performed a functional phenotyping of all recorded neuron using machine learning techniques in order to identify distinct neuronal subpopulations based on waveform and spiking features. We observed an 80% classification accuracy of neurons from TTAE mice in our logistic regression model and found that anatomy and not function is more predictive of exposure-induced deficits. We found that, despite HPF neurons sharing many physiological characteristics with thalamic (TH) neurons, TTAE has a bidirectional impact on the two populations, producing hypoexcitability in the TH and hyperexcitability in the HPF. This mirrors previous results found in Fos studies ^18^. Ultimately, we show that this approach provides a critical link between *ex vivo* observations and *in vivo* function, offering a comprehensive understanding of how developmental alcohol exposure impairs real-time circuit dynamics underlying spatial cognition.

## Methods

### Animals

All animal procedures were approved by the UNM-HSC Animal Care and Use Committee and were conducted in accordance with NIH guidelines. C57BL/6J breeding pairs were obtained from Jackson Labs and maintained on a 12-h reverse light/dark cycle (lights on at 8 p.m.). Animals were kept in micro-isolated systems with ventilated lid tops and cages (Lab Products, Seaford, DE). All animals had *ad libitum* access to water and rodent chow (Teklad 2920x, Envigo, Indianapolis, IN).

### Treatment

On PND 7, mice were subcutaneously injected twice with saline or ethanol (20% v/v in normal saline) at a dose of 2.5 g/kg spaced 2 hours apart, which results in peak blood ethanol concentrations nearing 400 and 500 mg/dL ^12^. Mice were returned to their litter until weaning on PND 20 and allowed to age at least until PND 60 (**Figure 1A**).

**Figure 1.**
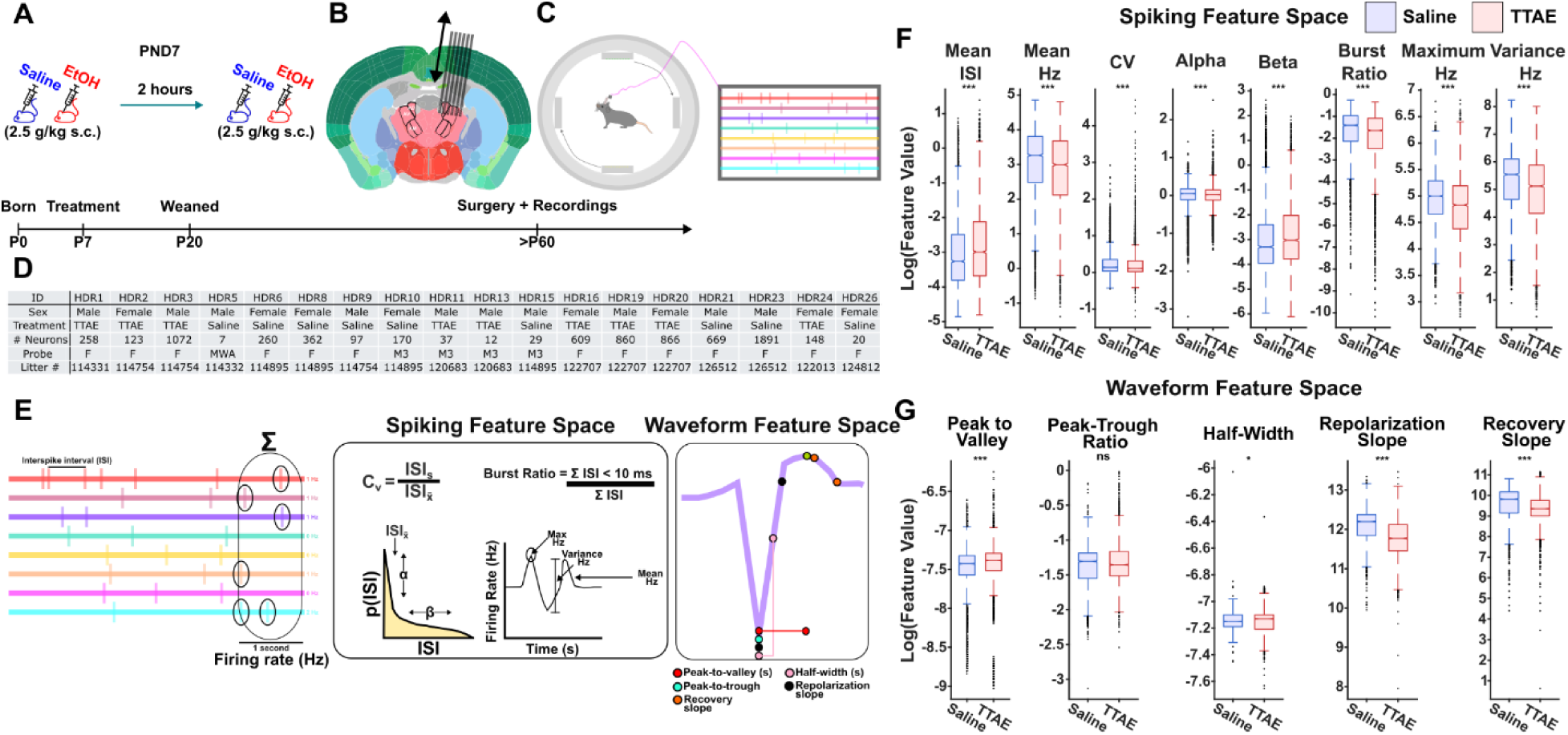
Experimental approach and effect of third-trimester equivalent alcohol exposure on spiking and waveform features. A. Illustration of the exposure paradigm. Mice on PND7 receive either saline or ethanol at a dose of 2.5 g/kg s.c. 2 hours are allowed to pass and mice receive a second dose of 2.5 g/kg s.c. of either saline or ethanol. B. On or above PND60, mice are implanted with an *in vivo* electrophysiology recording electrode. Most devices were on a movable microdrive that allowed lowering of the electrode after surgery. C. Mice were recorded from in a cylinder apparatus with dim LEDs on each quadrant of the walls. Only 2 LEDs were illuminated at any time 180 degrees from each other. The active LEDs rotated every 10 minutes. D. A table describing the treatment, sex, number of neurons, litter, and electrode type of each subject. Electrode type abbreviations are the following. **F** – indicates Cambridge Neurotech F-series silicon electrode. **MWA** – indicates multiwire array in-house fabricated device. **M3** - indicates ML-Precision M3 model silicon electrode. 96.6% of all neurons came from F devices on movable microdrives. 0.09% of all neurons came from fixed MWA. 3.3% came from fixed M3 devices. The sex and treatment breakdown are as follows: 5 male Saline (2693 neurons), 5 male TTAE (2239 neurons), 4 female Saline (812 neurons), and 4 female TTAE (1746 neurons). E. A description of the analyzed features, with the exception of features derived from autocorrelograms. F. Population-wide treatment differences are observed between saline and TTAE neurons. Broadly, the mean ISI and alpha parameter increase in TTAE neurons. The mean firing rate, CV, alpha parameter, burst ratio, maximum firing rate, and variance of the firing rate is decreased in TTAE neurons. G. Population-wide treatment differences are also observed in waveform characteristics. Broadly, the peak-to-valley time and half-width is increased in TTAE neurons. The repolarization slope and recovery slope are decreased in TTAE neurons.

### Surgery

On the day of surgery, adult mice were pulled from group housing and isolated. Following cessation of motor response, mice were pulled from the induction chamber and placed into the stereotaxic instrument. A rectangular craniotomy was performed medial to the AD (AP: 0.651, ML: 1.085, DV -3; Sunkin et al., 2013; **Figure 1B**). OptiBond (Kerr, USA) was applied to the skull surface and treated with blue light. Probes attached to 3D-printed moveable microdrives were then lowered at a 12-12.5-degree angle. Silicon elastomers (DuraGel, CambridgeNeurotech, UK; Kwik-Sil, WPI, USA) were applied to the open craniotomy to protect underlying tissue. Additional 3D-printed hardware was implanted to the skull to protect the electrode ^20^.

### Data acquisition and apparatus

Recordings began with the animal outside of the recording apparatus. After visual validation of both video and electrophysiology recordings, animals were placed inside the recording apparatus.

All hardware was controlled using Bonsai. All electrophysiology data was acquired utilizing an OpenEphys Acquisition Board (OpenEphys, Atlanta, GA). A Chameleon3 FLIR (Teledyne FLIR, UK) camera collected video at 40 FPS. Light-emitting diode (LED) cues were controlled using custom Arduino software. Animals were placed inside a 36.83-cm-diameter cylinder with gray walls. Affixed to the gray walls were 4 8×8 LED arrays arranged in 90-degree intervals. At any given time, only 2 180-degree opposite pairs of the 4 LED lights were active and displayed a blue X or green ^ across the full 8×8 array (similar to Duszkiewicz et al., 2024). LED arrays were pseudo randomly swapped on a roughly 9-minute inter-trial interval for a total of 2 hours. All LEDs were turned off 5 minutes before the end of the recording. All data were collected at 20 kHz and at broadband (1 – 6,000 Hz). Following daily data acquisition and the cessation of unique signal in that location, electrodes were lowered and allowed to settle overnight.

### Data preprocessing

Data were spikesorted using a combination of SpikeInterface and Kilosort4. Data were curated in SpikeInterface and MATLAB using quantitative spiking metrics, such as refractory violations and amplitude distributions. Videos were preprocessed in DeepLabCut and exported as CSV files. Broadband signals were downsampled to 1250 Hz for any LFP analyses.

### Histology and electrode verification

Post-hoc electrode verification was conducted using either serial sectioning or brain clearing. In serial sectioned tissue, electrode locations were tracked using DiI counterstained to DAPI and images were taken at 4x for both DAPI and DiI. Electrode locations were then marked in an Allen Brain Atlas CCFv3 3D volume using ITK-SNAP. Tissue underwent brain clearing following the CUBIC protocol and used CUBIC-standardized reagents (Tokyo Chemical Industry, CA, USA). Brains were imaged using a custom-built optical projection tomography system (https://github.com/AllenInstitute/AIBSOPT). Tomography was performed using NRecon (Bruker, MA, USA). Electrode tracts were manually traced in an ABA CCFv3 3D volume and exported to MATLAB. Custom software then used placement information, along with details on when and how much the electrode was lowered, to determine the placement of each individual neuron in the dataset. Additionally, placements that occurred in regions where spiking activity was not permissible, such as ventricles, were penalized and the broader placement was readjusted within 4 voxels (Supplementary Figure 1). The depth positions of our electrodes are shown in Supplementary Figure 3 and Supplementary Video 1.

### Feature curation and dimensionality reduction

Analyzed spiking features were with MATLAB. Mean waveforms were pulled for each neuron. Three-dimensional autocorrelograms (3D-ACGs) were generated to track features across neurons’ firing-rate deciles ^22^. 3D-ACGs were compressed to 2D and principal component analysis (PCA) was applied on both the extracted mean waveforms and 3D-ACGs. The full set of PCA components for both features was then utilized in a contrastive learning algorithm that minimized the distance between features of the same neuron in a high-dimensional space (1024D). This 1024D space was then reduced using UMAP. All algorithm parameters are listed in Supplementary Table 2. Broadly, our pipeline followed similar published pipelines, with the key difference being the use of PCA rather than an autoencoder for a first-pass dimensionality reduction ^22^.

### Clustering

Louvain clustering was used to further reduce the UMAP space into discrete clusters. Louvain parameters were selected in order to minimize the total number of clusters while maximizing the community coefficient. The number of clusters was minimized to ensure adequate sampling from both treatment conditions within each cluster. Parameters for clustering and UMAP-related analyses are listed in Supplementary Table 2.

### Statistics

#### Logistic regression

A general linear mixed-effects model was used in the present paper, with day, sex, and cluster as random effects. The least absolute shrinkage and selection operator (LASSO) was used to remove predictor variables that did not significantly contribute to the model. A cutoff of 1 standard error was used. Additionally, UMAP parameters were residualized in order to minimize the impact of treatment imbalances in the UMAP space. All predictor variables were z-score standardized prior to input into the model. Odds ratios (ORs) were univariate and reported in addition to receiver operating curve (ROC) characteristics and confusion matrices. An accurate saline classification was determined to be those below 0.5 and an accurate TTAE classification was determined to be those above 0.5. Our original logistic regression equation prior to LASSO is listed below as well as our residualization equations.

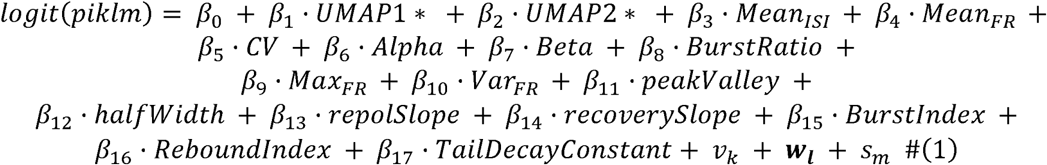

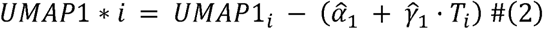

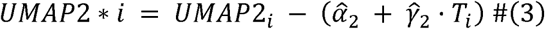

#### Pairwise comparisons

Generally, the features utilized were non-normally distributed, and non-parametric tests such as Wilcoxon’s Rank Sum were utilized where appropriate following log transformation. The resulting p-values were then FDR-corrected with an alpha of 0.05. Rose plots were generated by standardizing the entire feature population to the range [0, 1]. Subsequently, Spearman correlations were performed between standardized feature values. Treatment effect differences were calculated as a function of the difference between cluster Cohen’s d effect sizes.

## Results

Mice were exposed to saline or ethanol on PND7 (Figure 1A) and aged until adulthood. Electrodes were implanted that targeted the ATN at a 12–15-degree angle (Figure 1B). Following surgery recovery, mice were placed into our recording apparatus and recorded from (Figure 1C). Recordings were obtained from 18 animals in total (7490 neurons; Figure 1D). From our recorded neuronal population, we assessed both spike and waveform properties (Figure 1E). For each neuron, first order statistics related to spiking and waveforms were calculated in order to first determine if there are gross differences between treatment groups. We observed an increase in the mean interspike interval (ISI; Wilcoxon Rank Sum, p < 0.001, d = 0.119), a decrease in the mean firing frequency (Wilcoxon Rank Sum, p < 0.001, d = -0.119), a decrease in the coefficient of variation (CV; Wilcoxon Rank Sum, p < 0.001, d = -0.064), a decrease in the alpha parameter (Wilcoxon Rank Sum, p < 0.001, d = -0.050), an increase in the beta parameter (Wilcoxon Rank Sum, p < 0.001, d = 0.111), a decrease in the burst ratio (Wilcoxon Rank Sum, p < 0.001, d = -0.138), a decrease in the maximum firing frequency (Wilcoxon Rank Sum, p < 0.001, d = -0.174), and a decrease in firing frequency variance (Wilcoxon Rank Sum, p < 0.001, d = -0.180; Figure 1F). Waveform analysis revealed an increase in the peak-to-valley ratio (Wilcoxon Rank Sum, p < 0.001, d = 0.120), no effect on the peak-to-trough ratio (Wilcoxon Rank Sum, p = 0.107), a small decrease in the half-width of the waveform (Wilcoxon Rank Sum, p = 0.023, d = -0.030), and decreases in both the repolarization slope (Wilcoxon Rank Sum, p < 0.001, d = -0.417) and recovery slope (Wilcoxon Rank Sum, p < 0.001, d = -0.280; Figure 1G). Overall, our results suggest a broad decrease in excitability consistent with previous results. However, due to the size and heterogeneity of the recorded neuronal population, further analysis was required to delineate treatment effects on unique subpopulations.

Next, we applied UMAP to the 3D-ACGs and waveforms of each neuron in our dataset. Additional details found in Supplemental Figure 2. Partitioning of the UMAP embedding into 14 discrete clusters is shown in Figure 2A and B. Examination of the modal anatomical region within each cluster revealed that UMAP2 coordinates broadly align with a dorsal–ventral axis, whereas UMAP1 coordinates correspond more closely to a medial–lateral axis (Supplemental Figure 3). We then performed logistic regression with cluster, day, and sex as random effects. There was no systematic relationship between UMAP location and error rate (Figure 2C). We observed that increases in mean ISI were associated with a higher likelihood of TTAE classification whereas increases in the beta coefficient were once indicative of a higher likelihood of saline treatment (Figure 2D). Interestingly, the model had differing rates of false-positives versus false-negatives depending on the cluster examined. For instance, cluster 2 – which represents our HPF neurons – had the highest rate of saline neurons that were labeled as TTAE neurons (Figure 2E). Despite this, the majority of our clusters had greater than a 70% accuracy (Figure 2F). Our overall classification accuracy for both saline and TTAE conditions hovered at or near 80% (Figure 2G). Receiver operating characteristic (ROC) curve analysis indicated an overall model accuracy of 0.864 (Figure 2H).

**Figure 2.**
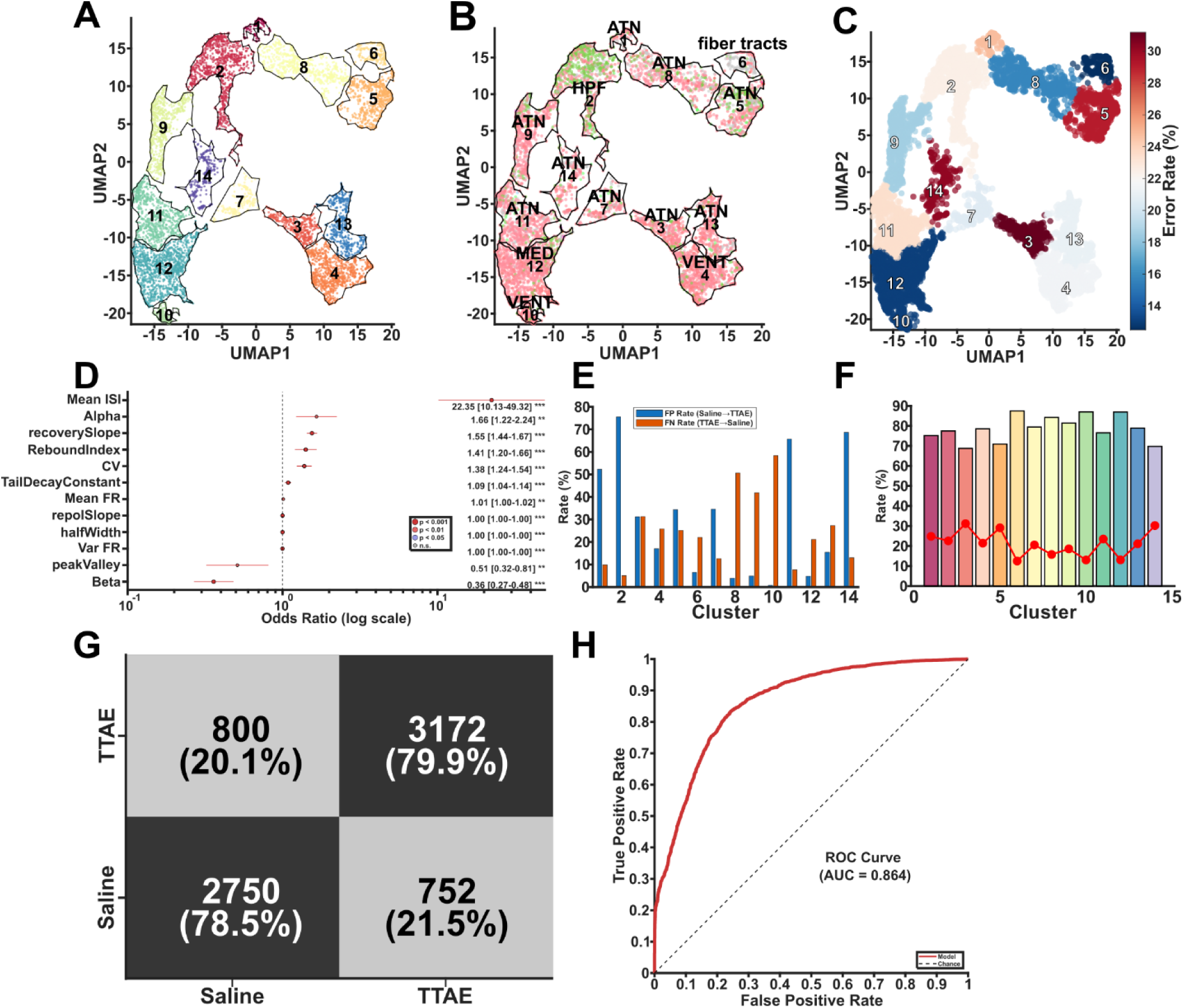
Logistic Regression Classification of Saline and TTAE Neurons Using UMAP-Embedded Features. A. The UMAP reduction was further reduced into 14 unique clusters using Louvain clustering. B. Dominant anatomical region within each cluster. For each of the 14 Louvain-defined clusters, the most frequent anatomical region label is displayed. This mapping reveals that UMAP2 coordinates broadly align with a dorsal–ventral axis, while UMAP1 coordinates correspond to a medial–lateral axis, indicating that the UMAP embedding captures aspects of anatomical organization (need to define ATN, VENT, HPF, and MED abbreviations). C. The classification error rate is shown for each cluster. D. Increases in the mean ISI is highly predictive of TTAE while increases in beta is the main predictor of saline treatment. Additional features are discriminative. Values to the left of the midline predict saline treatment; values to the right predict TTAE. E. The false-positive and false-negative rates are shown for each cluster. F. The accuracy and error rate for each cluster is shown. G. TTAE neurons were classified accurately 79.9% of the time. Saline neurons were improved to 78.5% accuracy. H. The receiver operating characteristic indicates that the model has good sensitivity for predicting TTAE neurons.

The unequal distribution of error rates in our cluster model invited further exploration. Figure 3A shows a sample of the critical features our logistic regression model identified, overlaid on our UMAP reduction. In these examples, the peakValley ratio is markedly lower in clusters predominantly labeled as fiber tracts, whereas features such as mean ISI and the beta coefficient display broader, globally distributed differences. We then performed pairwise non-parametric comparisons across each cluster-feature condition (Figure 3B). While many of our clusters are similar in their treatment effects, outliers such as cluster 2 were observed to increase in their excitability. Additionally, cluster 2 was predominately labeled as comprising HPF neurons (Figure 2B) indicating a differential impact from TTAE. Next, we summed the absolute value of each Cohen’s D for each feature tested to determine which features were most impacted by our treatment (Figure 3C). From this, we conclude that the repolarization slope and recovery slope are both highly impacted by TTAE. Additionally, the burst ratio, which reflects the propensity of neurons to fire in bursts, was strongly affected. A full table of pairwise comparisons can be found in Supplemental Table 1. Together, these features provide a basis for generating further hypotheses regarding the functional impact of TTAE.

**Figure 3.**
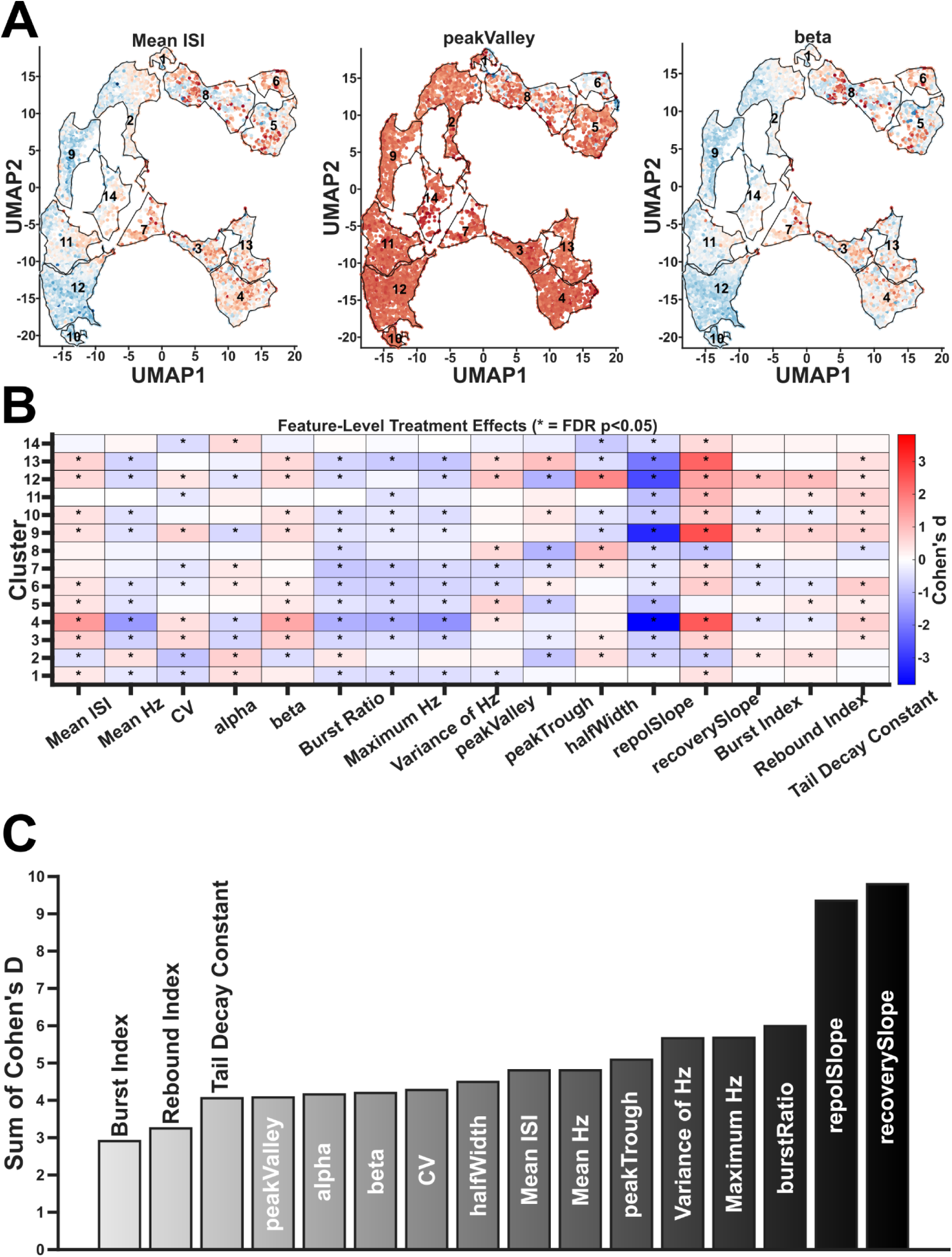
Pairwise comparisons of spike and waveform features within a cluster. A. Example spike and waveform characteristics overlaid on the UMAP reduction. Redder values indicate a higher value. Bluer values indicate a lower value. B. Pairwise comparisons of each feature within a cluster. Importantly, cluster 2 is higher in firing rate, all other clusters are either null or lower in firing rate. Asterisks indicate significant differences in features between saline and TTAE that survived FDR correction. C. The absolute sum of the Cohen’s D value is reported. The recovery slope, repolarization slope, and burst ratio are the top three features most impacted by TTAE.

To further investigate the effect of TTAE, a collection of rose plots were plotted for the saline and TTAE conditions of each cluster in order to visualize which clusters have physiological similarities (Figure 4A). We then performed a Spearman correlation on the collection of physiological features in our rose plots. We observe, for example, large correlations between clusters high in excitability such as 9 and 12 (Figure 4B). Last, we determined if similarities in cluster physiology were correlated with treatment effects. In order to determine whether similarities in physiology result in similarities in treatment effects, we correlated the pairwise difference in Cohen’s D effect sizes with the pairwise R-value from each cluster. We observed a small but significant correlation, suggesting that physiologically similar clusters generally experience a similar impact from TTAE (Figure 4C). However, notable exceptions to this trend were observed in the top right of the figure, suggesting that some highly correlated clusters are not experiencing the same treatment effect. To further refine this point, we constructed network graphs of both our physiological and treatment effect correlations (Figure 4D,E). In our physiology correlation network, we observe two modules that generally delineate along the lines of overall excitability (Figure 4D). However, the treatment correlation network reveals four modules of treatment effects (Figure 4E). This suggests that anatomically distinct regions differentiate TTAE effects more strongly than overlapping physiological features.

**Figure 4.**
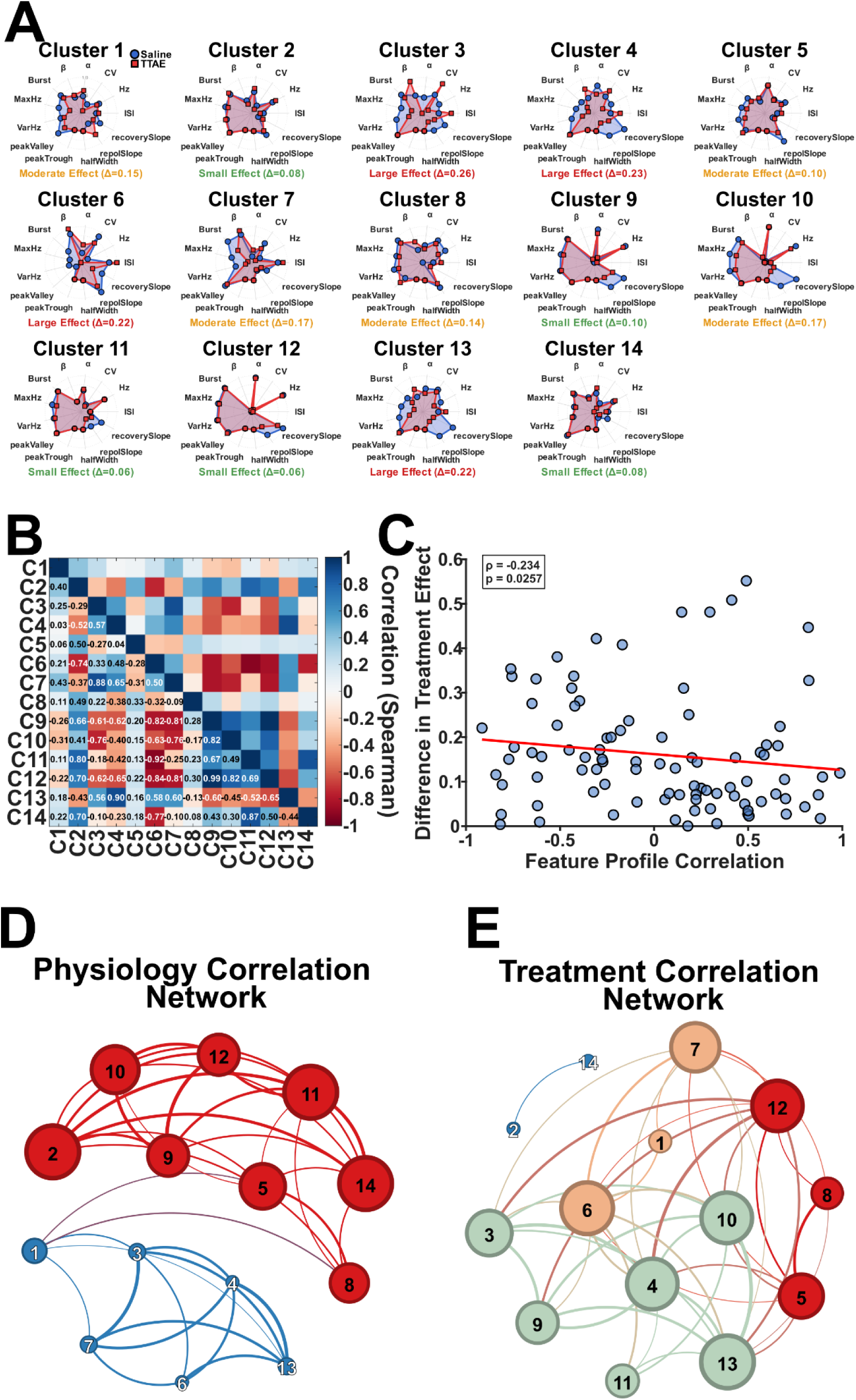
Physiological feature profiling of each cluster. A. Rose plots were generated for each cluster of each feature. The features were standardized to the range [0, 1] across the full population, such that 1 indicates the highest value in the dataset and 0 the lowest. B. Spearman correlations were performed between all clusters with all features as the correlates. C. The correlation values from B are plotted against the difference between Cohen’s D values between each cluster. D. Two large physiology correlation modules emerge (each represented by red or blue colors). E. Despite two large physiology correlation values emerging, 4 modules related to treatment effects emerge (represented by orange, red, green, and blue colors), suggesting that overlaps in physiology are not necessarily predictive of TTAE impacts.

Last, we directly compared our HPF population of neurons to our TH population of neurons. Our TH condition had 2772 and 2836 in the saline and TTAE conditions, respectively.

Our HPF condition had 249 and 597 neurons in the saline and TTAE conditions, respectively. Pairwise comparisons were performed between saline and TTAE neurons in their respective TH and HPF brain regions. HPF neurons have a general increase in excitability whereas TH neurons have a general decrease in excitability (Figure 5A). Next, we performed PCA in order to reduce our spike and waveform space into 2 principal components representing excitability and waveforms and bursting (Figure 5B). Following this, we plotted the centroid for each treatment and brain region combination and the distance they move in PCA space. Interestingly, we observe that TH neurons shift leftward along PC1, indicating a decrease in overall excitability. HPF neurons are shifted rightward on PC1, indicating an increase in overall excitability (Figure 5C). Last, based on our overall decreases in TH excitability and increases in HPF excitability, we posit a conceptual model in Figure 5D in which information fidelity may be lost due to TH hypoexcitability, potentially compensated by HPF hyperexcitability.

**Figure 5.**
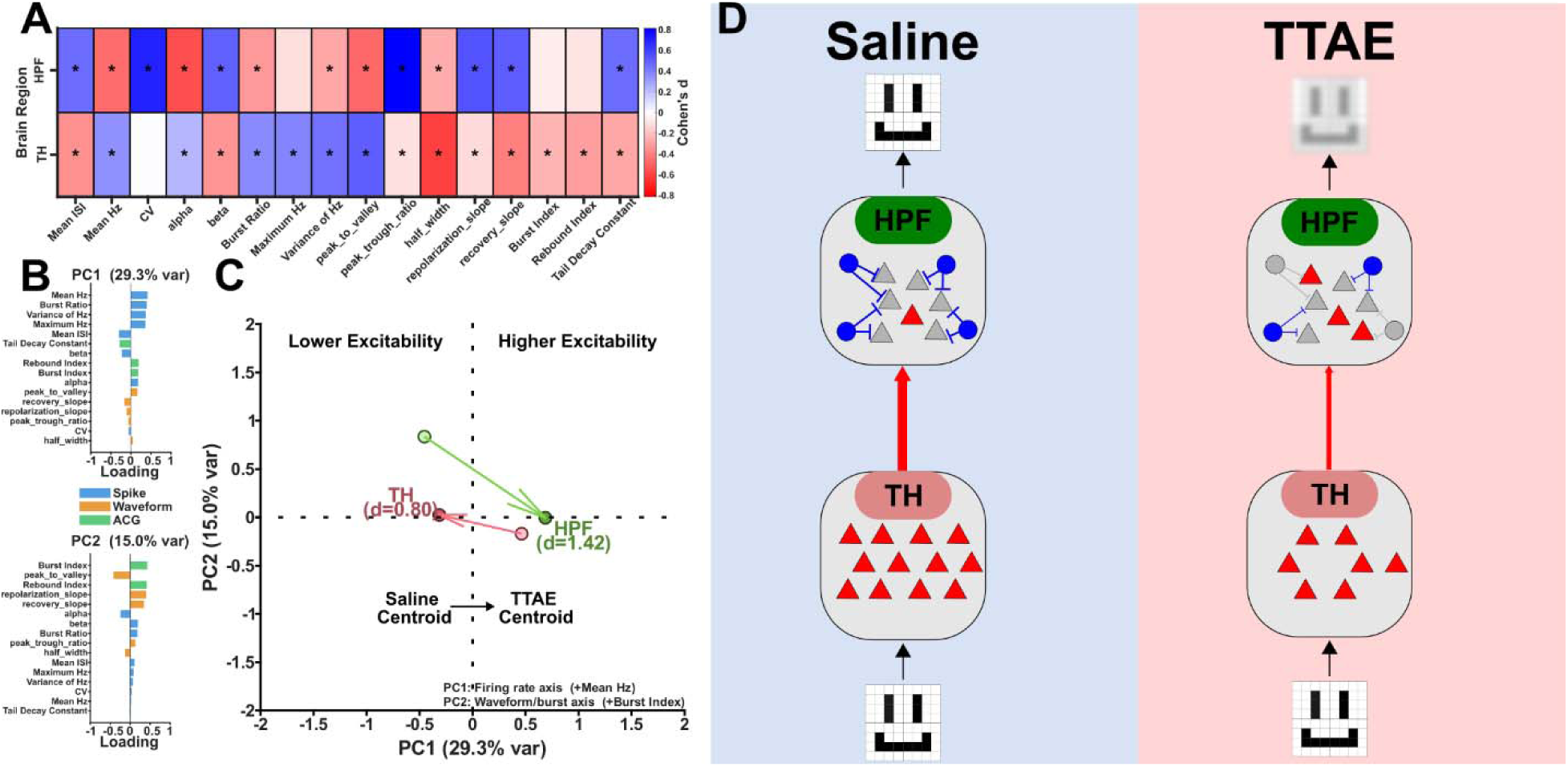
Direct Thalamus-Hippocampus comparison. A. Neurons were assigned to either TH and HPF conditions based on their histology. Our TH condition had 2772 and 2836 in the saline and TTAE conditions, respectively. Our HPF condition had 249 and 597 neurons in the saline and TTAE conditions, respectively. Each spike and waveform feature was tested pairwise between saline and TTAE conditions in both brain regions. Cohen’s d shading indicates the effect size of each comparison, with red indicating values higher in TTAE. Asterisks indicate significant pairwise comparisons that survived FDR corrections. B. Next, principal component analysis was performed on all spike and waveform features. The principal component loadings for the first two PCs and their respective features are depicted. PC1 is predominantly an excitability axis. PC2 is predominantly a waveform- and burst-feature axis. C. The centroid is depicted for both TH and HPF data points under the saline and TTAE conditions. The graph indicates shifts in the state space following TTAE. The arrow points from saline to TTAE. Critically, we observe a rightward shift in PC1 for HPF neurons and a leftward shift in PC1 for TH neurons, indicating bidirectional effects of TTAE exposure. The distance value (d) is plotted. D. Conceptual model of TTAE-induced circuit adaptations in thalamic and hippocampal neurons. TTAE decreases the excitability of thalamic (TH) glutamatergic neurons (red triangles), while hippocampal (HPF) pyramidal neurons (red and gray triangles) exhibit increased excitability, likely driven by a reduction in local interneuron activity (blue and gray circles). This bidirectional modulation of excitability may reduce information fidelity across circuits, contributing to the learning and memory deficits observed in FASD.

## Discussion

The in vivo functional consequences of TTAE remain strikingly underexplored, and this study represents the first large-scale profiling of its effects in freely behaving mice. While many hippocampal and thalamic neurons exhibited overlapping functional phenotypes, hippocampal neurons showed increased excitability, whereas thalamic neurons showed decreased excitability. These findings reveal region-specific circuit adaptations to developmental alcohol exposure and highlight the functional diversity of thousands of neurons across anatomically distinct zones. By linking anatomical specificity to functional outcomes, this work offers a compelling and innovative framework for understanding how early-life alcohol exposure reshapes neural circuits, while also informing the development of targeted interventions for FASDs.

The present results are in general agreement with previously published work showing increases in hippocampal c-Fos expression and decreases in the ATN following contextual fear conditioning. Our conceptual model posits that decreases in TH excitability are compensated for by increases in HPF excitability. Mechanistically, this increase in excitability may be due to decreases in HPF interneurons following TTAE ^23,24^. Additionally, TTAE has been shown to trigger apoptotic neurodegeneration in cortical interneurons associated with a compensatory increase in their function in areas such as the RSC ^25^. Taken together, this suggests that broad reductions in TH excitability may be compensated for by increases in cortical and HPF excitability. The tradeoff of this compensation may be deficits in information fidelity, leading to reductions in learning and memory observed following both TTAE and prenatal alcohol exposure (PAE). Deficits in neuronal excitability could subsequently lead to difficulties in stabilizing neuronal representations, such as hippocampal head-direction and place cells, and in learning and memory dependent on the hippocampus ^26–28^. Indeed, PAE has been shown to cause deficits in hippocampal place cell firing as well as theta rhythmicity that may be related to the neuron’s excitability ^29^. The clinical impacts of these decreases in excitability may include deficits in spatial cognition observed in children with FASD ^30,31^.

Both the repolarization and recovery slopes were significantly affected in the majority of our clusters, resulting in high overall Cohen’s D values. Generally, the repolarization slope was higher in saline-exposed neurons and lower in TTAE neurons, indicating a slower repolarization following action potentials in TTAE neurons, which may explain some of the hypoexcitability observed. Interestingly, the HPF-associated cluster also showed a slower repolarization slope, indicating that the TTAE impacts on individual neurons may be similar, but differences in microcircuitry produce the observed hyperexcitability phenotype. Voltage-gated sodium channels and voltage-gated potassium channels contribute to the repolarization slope, and at the time of writing, two studies have investigated voltage-gated sodium channels following developmental alcohol exposure and have found that they are disrupted both functionally and transcriptionally ^32,33^. Ultimately, this suggests that the voltage-gated sodium channels may be involved in the observed excitability phenotypes.

A key feature of interest that showed significant differences between treatment groups was the burst ratio. Thalamic cells show a high degree of bursting onto cortical cells, likely as a mechanism for regulating perceptual and attentional processes, and have been observed across multiple species and thalamic nuclei ^34,35^. Mechanisms that regulate bursting, such as the hyperpolarization-activated current (I_h_) may be key to understanding our broad decreases in excitability. Furthermore, recent work has shown that *Hcn1*, the gene that encodes the HCN1 channel that partially contributes to I_h_ currents, is highly and differentially expressed in the AD relative to its neighboring thalamic subnuclei ^36^. Investigating it and other differentially expressed genes in the AD, such as *Chrm1*^36^, may further elucidate TTAE-related disruptions.

The accuracy of the logistic regression indicates that we can make meaningful inferences about a neuron’s treatment status based on spike and waveform features alone. Extrapolating from the level of individual neurons, we should observe differences in aggregate electrophysiological signals, such as local field potentials and electroencephalograms , given that TH populations show hypoactivity and HPF show hyperactivity. Clinically, this may be valuable in helping diagnose those with FASDs. Additionally, the alpha band may inform us about corticothalamic communication in rodent models and has been linked to deficits in FASD ^37–39^. Robust phenotyping of electroencephalogram signals in PAE rodents and, subsequently, in humans may critically aid diagnosis.

Recent work has demonstrated that aggregate electrophysiological features can be used to classify and cluster large neuronal populations. Our approach was inspired by the IBL-NEMO framework, which has shown moderate success in large-scale datasets containing thousands of neurons when classifying anatomical regions or interneuron subtypes ^22,40^.The same framework has also been applied to smaller datasets of optically evoked cerebellar neurons, where spike and waveform features enabled accurate classification of cerebellar subtypes ^40^. Despite the relative success of the present paper, classification of neuronal subtypes based on electrophysiological features remains an active field of research that will continue to improve.

The present study has several key limitations worth highlighting. First, there is no ground truth in this dataset. Previous studies have paired optogenetics with *in vivo* electrophysiology to isolate genetically defined subpopulations of neurons, which can serve as ground truth datasets ^40^, a strategy we could adopt in future studies. Although histology confirmed the general trajectory of each electrode track, the precise probe location after each lowering step could only be approximated. Consequently, we referenced broader hierarchical regions from the Allen Brain Atlas rather than assigning neurons to specific thalamic nuclei. Future work will combine histological reconstruction with electrophysiological signatures from both spikes and LFPs to more precisely localize recordings and resolve activity within individual thalamic subnuclei.

Overall, using UMAP enabled functional segregation of neural populations and moderately accurate classification of TTAE-exposed neurons. Across the recorded population, we observed bidirectional changes in excitability, with hippocampal neurons exhibiting increased activity while thalamic neurons were generally suppressed. These findings indicate that the effects of developmental alcohol exposure are strongly shaped by anatomical context. Importantly, this study introduces a novel analytical framework to the FASD field, combining large-scale *in vivo* electrophysiological recordings in freely moving animals with dimensionality reduction and machine-learning–based classification to characterize neuronal function at the population level. By capturing the activity of thousands of neurons across anatomically distinct regions, this approach provides a powerful strategy for identifying circuit-level adaptations following developmental alcohol exposure. Together, these results reveal regionally divergent circuit responses to TTAE and establish a scalable methodological framework for further dissecting the complex neural mechanisms underlying cognitive deficits associated with FASD.

## Author Contributions

MDM and GA performed all data collection. Analysis was led by MDM, DNL, BJC and SAM. Conceptualization was performed by CFV, BJC, DNL, and MDM.

## References

1. Dobbing, J. & Sands, J. Comparative aspects of the brain growth spurt. Early Hum. Dev. 3, 79–83 (1979).

2. Umer, A. et al. Prevalence of alcohol use in late pregnancy. Pediatr. Res. 88, 312–319 (2020).

3. Bubb, E. J., Kinnavane, L. & Aggleton, J. P. Hippocampal - diencephalic - cingulate networks for memory and emotion: An anatomical guide. Brain Neurosci. Adv. 1, 2398212817723443 (2017).

4. Papez, J. W. A proposed mechanism of emotion. Arch. Neurol. Psychiatry 38, 725–743 (1937).

5. Vogt, B. A. Cingulate cortex in the three limbic subsystems. Handb. Clin. Neurol. 166, 39–51 (2019).

6. Clark, B. J. & Harvey, R. E. Do the anterior and lateral thalamic nuclei make distinct contributions to spatial representation and memory? Neurobiol. Learn. Mem. 133, 69–78 (2016).

7. Taube, J. S. The head direction signal: origins and sensory-motor integration. Annu. Rev. Neurosci. 30, 181–207 (2007).

8. Clark, B. J., Bassett, J. P., Wang, S. S. & Taube, J. S. Impaired Head Direction Cell Representation in the Anterodorsal Thalamus after Lesions of the Retrosplenial Cortex. J. Neurosci. 30, 5289–5302 (2010).

9. Roy, D. S. et al. Anterior thalamic dysfunction underlies cognitive deficits in a subset of neuropsychiatric disease models. Neuron 109, 2590–2603.e13 (2021).

10. Vantomme, G. et al. A Thalamic Reticular Circuit for Head Direction Cell Tuning and Spatial Navigation. Cell Rep. 31, 107747 (2020).

11. Ikonomidou, C. et al. Ethanol-induced apoptotic neurodegeneration and fetal alcohol syndrome. Science 287, 1056–1060 (2000).

12. Wozniak, D. et al. Apoptotic neurodegeneration induced by ethanol in neonatal mice is associated with profound learning/memory deficits in juveniles followed by progressive functional recovery in adults. Neurobiol. Dis. 17, 403–414 (2004).

13. Harvey, R. E., Berkowitz, L. E., Hamilton, D. A. & Clark, B. J. The effects of developmental alcohol exposure on the neurobiology of spatial processing. Neurosci. Biobehav. Rev. 107, 775–794 (2019).

14. Woods, K. J. et al. Prenatal alcohol exposure affects brain function during place learning in a virtual environment differently in boys and girls. Brain Behav. 8, e01103 (2018).

15. Granato, A., Santarelli, M., Sbriccoli, A. & Minciacchi, D. Multifaceted alterations of the thalamo-cortico-thalamic loop in adult rats prenatally exposed to ethanol. Anat. Embryol. (Berl.) 191, 11–23 (1995).

16. Rodriguez, C. I., Davies, S., Calhoun, V., Savage, D. D. & Hamilton, D. A. Moderate Prenatal Alcohol Exposure Alters Functional Connectivity in the Adult Rat Brain. Alcohol. Clin. Exp. Res. 40, 2134–2146 (2016).

17. Bird, C. W. et al. Binge-like ethanol exposure during the brain growth spurt disrupts the function of retrosplenial cortex-projecting anterior thalamic neurons in adolescent mice. Neuropharmacology 241, 109738 (2023).

18. Morningstar, M. D. et al. Connectivity of the neuronal network for contextual fear memory is disrupted in a mouse model of third-trimester binge-like ethanol exposure. Alcohol Clin. Exp. Res. 49, 315–331 (2025).

19. Sunkin, S. M. et al. Allen Brain Atlas: an integrated spatio-temporal portal for exploring the central nervous system. Nucleic Acids Res. 41, D996–D1008 (2013).

20. Vöröslakos, M., et al. 3D-printed Recoverable Microdrive and Base Plate System for Rodent Electrophysiology. Bio-Protoc. 11, e4137 (2021).

21. Duszkiewicz, A. J. et al. Local origin of excitatory–inhibitory tuning equivalence in a cortical network. Nat. Neurosci. 27, 782–792 (2024).

22. Yu, H. et al. In vivo cell-type and brain region classification via multimodal contrastive learning. in (2024).

23. Bird, C. W., Taylor, D. H., Pinkowski, N. J., Chavez, G. J. & Valenzuela, C. F. Long-term Reductions in the Population of GABAergic Interneurons in the Mouse Hippocampus following Developmental Ethanol Exposure. Neuroscience 383, 60–73 (2018).

24. Lopez, K. M. et al. Impact of acute binge-like ethanol exposure during the third-trimester equivalent on subicular interneurons in mice. Am. J. Drug Alcohol Abuse 51, 717–729 (2025).

25. Bird, C. W., Chavez, G. J., Barber, M. J. & Valenzuela, C. F. Enhancement of parvalbumin interneuron-mediated neurotransmission in the retrosplenial cortex of adolescent mice following third trimester-equivalent ethanol exposure. Sci. Rep. 11, 1716 (2021).

26. Diersch, N., Valdes-Herrera, J. P., Tempelmann, C. & Wolbers, T. Increased Hippocampal Excitability and Altered Learning Dynamics Mediate Cognitive Mapping Deficits in Human Aging. J. Neurosci. Off. J. Soc. Neurosci. 41, 3204–3221 (2021).

27. Reagh, Z. M. et al. Functional Imbalance of Anterolateral Entorhinal Cortex and Hippocampal Dentate/CA3 Underlies Age-Related Object Pattern Separation Deficits. Neuron 97, 1187–1198.e4 (2018).

28. Yassa, M. A. et al. Pattern separation deficits associated with increased hippocampal CA3 and dentate gyrus activity in nondemented older adults. Hippocampus 21, 968–979 (2011).

29. Harvey, R. E., Berkowitz, L. E., Savage, D. D., Hamilton, D. A. & Clark, B. J. Altered Hippocampal Place Cell Representation and Theta Rhythmicity following Moderate Prenatal Alcohol Exposure. Curr. Biol. CB 30, 3556–3569.e5 (2020).

30. Dodge, N. C. et al. Reduced Hippocampal Volumes Partially Mediate Effects of Prenatal Alcohol Exposure on Spatial Navigation on a Virtual Water Maze Task in Children. Alcohol. Clin. Exp. Res. 44, 844–855 (2020).

31. Willoughby, K. A., Sheard, E. D., Nash, K. & Rovet, J. Effects of prenatal alcohol exposure on hippocampal volume, verbal learning, and verbal and spatial recall in late childhood. J. Int. Neuropsychol. Soc. JINS 14, 1022–1033 (2008).

32. Chatterjee, D., Mahabir, S., Chatterjee, D. & Gerlai, R. Lasting effects of mild embryonic ethanol exposure on voltage-gated ion channels in adult zebrafish brain. Prog. Neuropsychopharmacol. Biol. Psychiatry 110, 110327 (2021).

33. Xu, W. et al. Genome-Wide Transcriptome Landscape of Embryonic Brain-Derived Neural Stem Cells Exposed to Alcohol with Strain-Specific Cross-Examination in BL6 and CD1 Mice. Sci. Rep. 9, 206 (2019).

34. Lesica, N. A. & Stanley, G. B. Encoding of Natural Scene Movies by Tonic and Burst Spikes in the Lateral Geniculate Nucleus. J. Neurosci. 24, 10731–10740 (2004).

35. Swadlow, H. A. & Gusev, A. G. The impact of ‘bursting’ thalamic impulses at a neocortical synapse. Nat. Neurosci. 4, 402–408 (2001).

36. Kapustina, M. et al. The cell-type-specific spatial organization of the anterior thalamic nuclei of the mouse brain. Cell Rep. 43, 113842 (2024).

37. Andrew, C., Meintjes, E., Jacobson, S., Jacobson, J. & Molteno, C. Resting and task-related alpha-band EEG oscillations differentiate children diagnosed with Fetal Alcohol Spectrum Disorder. NeuroImage 47, S70 (2009).

38. da Silva, F. H., van Lierop, T. H., Schrijer, C. F. & van Leeuwen, W. S. Organization of thalamic and cortical alpha rhythms: spectra and coherences. Electroencephalogr. Clin. Neurophysiol. 35, 627–639 (1973).

39. Klimesch, W. Alpha-band oscillations, attention, and controlled access to stored information. Trends Cogn. Sci. 16, 606–617 (2012).

40. Beau, M. et al. A deep learning strategy to identify cell types across species from high-density extracellular recordings. Cell 188, 2218–2234.e22 (2025).

