## Supplemental Figures 1-3, Supplemental Tables 1-2 for "Large-Scale In Vivo Electrophysiology and Dimensionality-Reduction Analysis Reveal Opposing Thalamic and Hippocampal Excitability After Third-Trimester–Equivalent Alcohol Exposure in Mice"

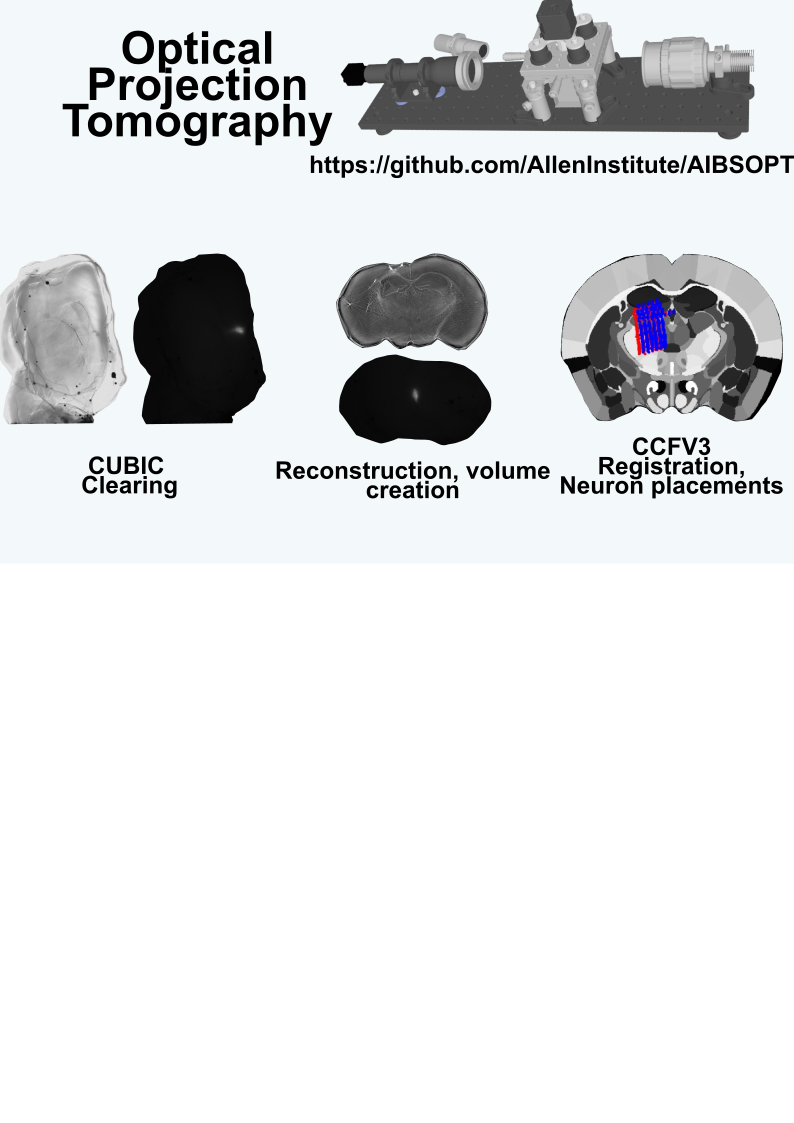

Figure 1

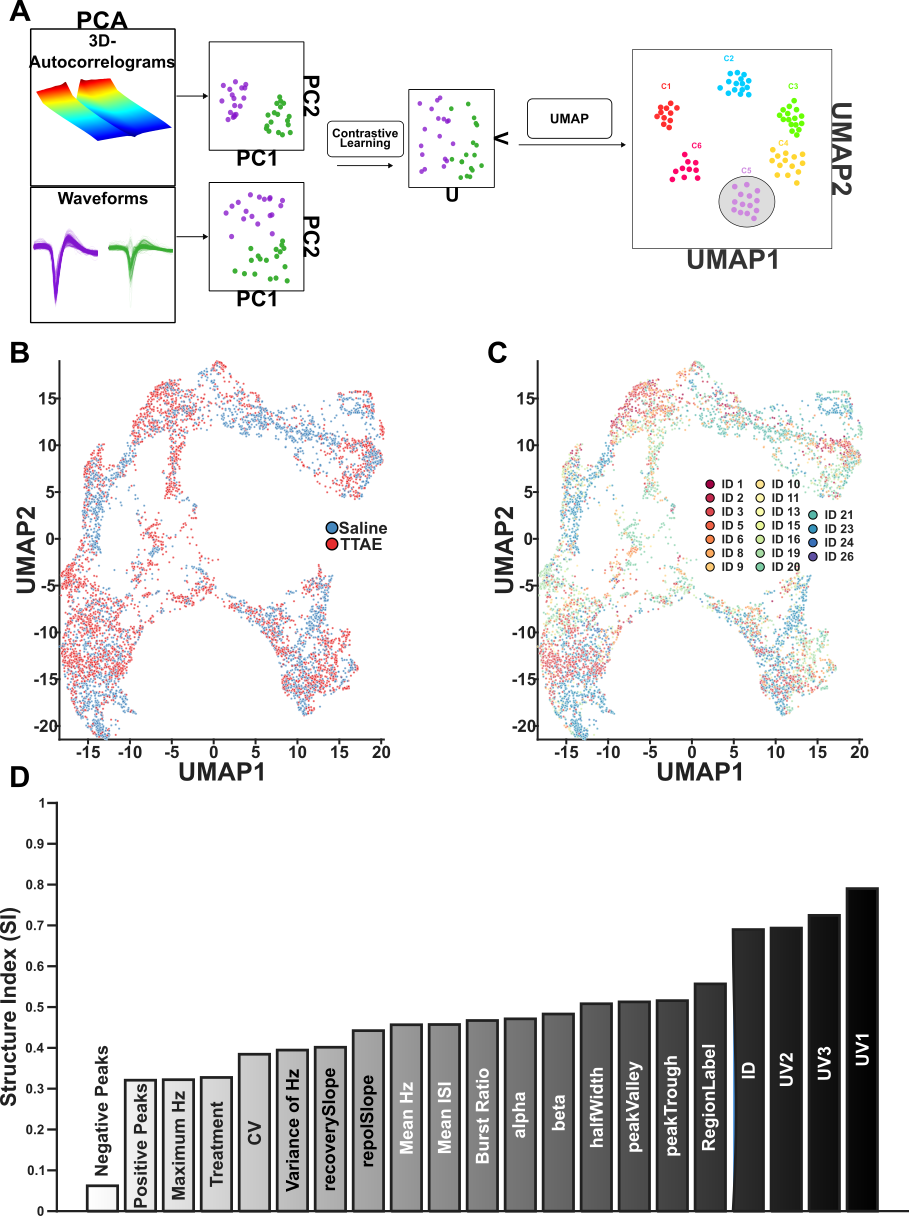

Figure 2

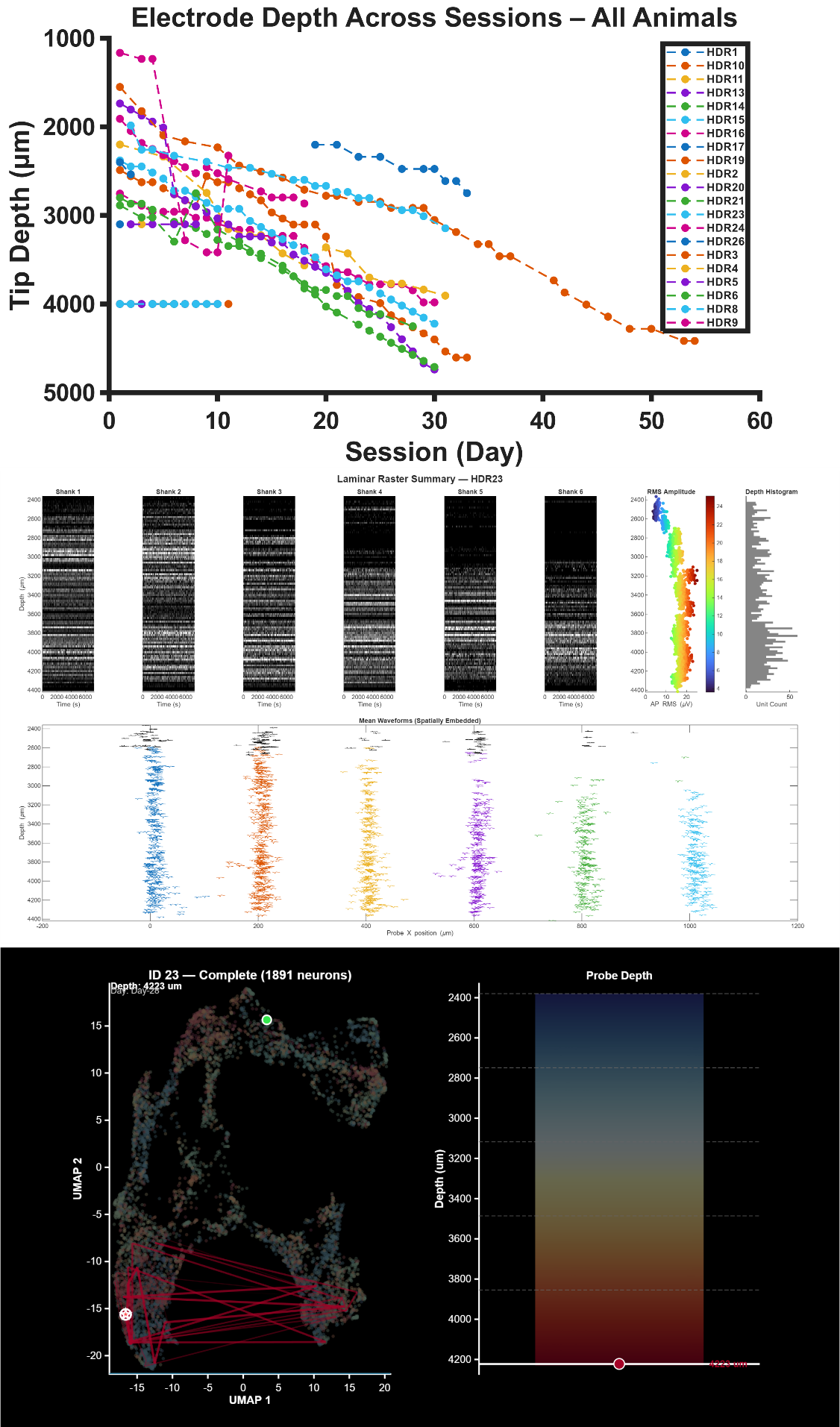

Figure 3

Table 1

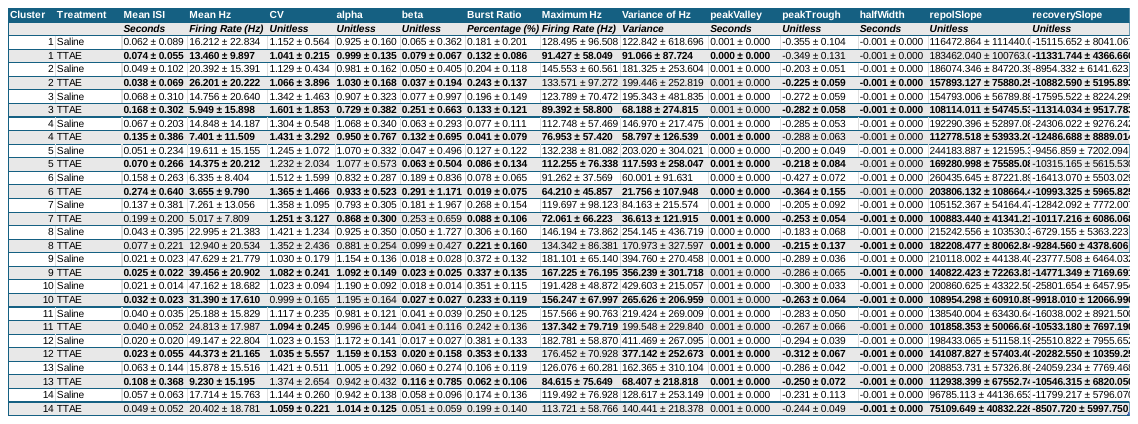

Table 2

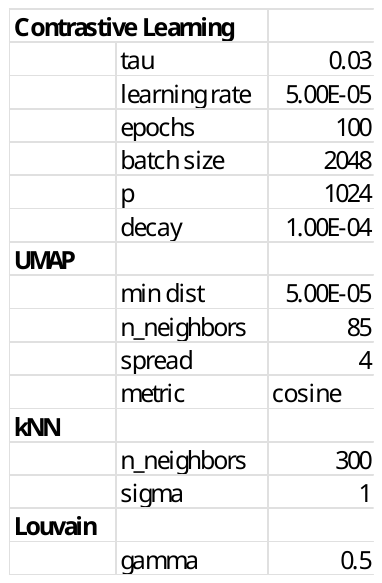

**Supplementary Figure and Table Captions**

**Figure 1. Optical projection tomography.** An optical projection tomographer (OPT) was constructed following opensource designs provided by the Allen Brain Institute. Broadly, a cleared brain is placed on a rotating disc. Light is uniformly shown through the disk through a diffuser lens. Fluorescence is excited using an LED of appropriate wavelength. 400 images are taken at a 0.9 angle. The resulting images are processed using tomography software. In our case, we used NRecon. This results in serially sectioned images from the full brain. Next, a volume was constructed using similar standards and meta data as the Allen Brain Common Coordinate Framework at 25 micron voxels. Following volume construction, placement of fluorescent tracts were labeled in ITK-SNAP inside of a standard ABA CCFV3 25 micron volume. The tracts of the probe were reconstructed using the record of electrode turns over each session. Neuron spatial coordinates were obtained as a result of the spikesorting process. The spatial coordinates were transformed using the depth information and put into ABA CCFV3 space. Neurons then had a spatial voxel assigned to them. Each spatial voxel has its own brain region in Allen Brain space. Neurons then were assigned their relevant brain region.

**Figure 2. UMAP-Based Embedding of Electrophysiological Features Reveals Structured Neuronal Populations.** A. Workflow for UMAP-based dimensionality reduction of electrophysiological features. First, 3D-ACGs and waveforms underwent principal component analysis. The full set of principal components were then utilized in a contrastive learning algorithm to embed the features of the same neuron in a common space. UMAP was then performed on the results of the contrastive learning algorithm. B. The UMAP reduction is depicted with saline and TTAE neurons overlaid. There is a general balance in where each treatment is represented in the UMAP reduction. C. The UMAP reduction is depicted with the animal ID overlaid. Importantly, as the majority of our neurons were recorded from animals with microdrives, the overlaid ID depicts the location of each recorded population as the drive is lowered. D. The structure index (SI) was calculated. The top three dimensions from our contrastive learning and ID of the animal explain the most structure of the UMAP reduction.

**Figure 3. Depth is recapitulated inside of UMAP space.** The depth per session is shown. Generally each probe follows a similar tract and covers similar positions varying mostly in where they start and ultimately end. Generally, we sought to record from 30 sessions per animal. Next, the laminar distribution from a single animal is shown. Spike rates are shown on top as a function of depth. Waveforms are shown on bottom as a function of depth. The waveforms on bottom are colored according to the shank they belong to. Waveforms in black are upward deflecting and presumed to be from fiber tracts. Thus, in this particular animal, we start in the fiber tracts immediately above the thalamus and move our way through the thalamus. Last, the UMAP is shown. Overlaid on the UMAP in green is the first neuron from the highest depth in a single subject. As depth is increased, the distribution of neurons is lowered across UMAP2 suggesting that UMAP2 is tracking depth. UMAP1 seems to be tracking the medial-lateral position of each neuron.

**Table 1.** Each pairwise comparison is provided. Bolded values in the TTAE row indicate a significant difference between saline and TTAE animals.

**Table 2.** Each parameter used across our analyses is shown.
